# Structure of Connexin26 from *Lepidosiren paradoxa*

**DOI:** 10.64898/2026.02.11.705270

**Authors:** Deborah H. Brotherton, Nicholas Dale, Alexander D. Cameron

## Abstract

Connexin gap junction channels enable the direct exchange of molecules and ions between cells. Channel opening is regulated by various physiological stimuli. Human connexin26 gap junction channels close in response to elevated levels of CO_2_ in a process that is independent of pH, with structures demonstrating a [CO_2_]-dependent conformational change. Cx26 from the lungfish, *Lepidosiren paradoxa* is also CO_2_ sensitive. Here we solve its structure at high [CO_2_]. We observe an open conformation of the protein where the N-terminal helix that influences the aperture of the pore is pulled away from the centre. This conformation is stabilised by the presence of a detergent binding between TM3 and TM4 that would prevent the conformational changes necessary to close the protein upon exchange to high [CO_2_]. The structure supports a mechanism in which the conformation of a motif shown to be important for CO_2_ sensitivity is correlated with the opening of the pore.

## Introduction

Connexins form large pore channels that allow the exchange of molecules and ions either with the extracellular milieu, in the form of hexameric hemichannels (connexons) or with neighbouring cells when hemichannels dock to form gap junction channels (GJCs) ^1^. There are 21 connexin genes grouped into different classes ^2^. Mutations in connexins lead to severe diseases ^3^. In connexin26 (Cx26), for example, mutations are a leading cause of congenital deafness and can lead to keratitis ichthyosis deafness syndrome ^4^, while X-linked Charcot Marie Tooth disease is caused by the loss of function of Cx32 ^5,6^. Connexins have been shown to open or close in response to various stimuli, including voltage ^7,8^, pH ^9–11^, or intracellular Ca^2+^ concentrations ^12^. One well known stimulus is the response to the partial pressure of CO_2_ (PCO_2_). This was thought to be purely an indirect effect due to PCO_2_ modulating the pH. However, we have observed that a subset of connexins from the β and α families, respond to CO_2_ independently of pH ^13–17^. For these connexins, hemichannels are opened by increasing PCO_2_ whereas they are closed by decreasing pH. While Cx26, Cx30, Cx32, Cx43 and Cx50 hemichannels respond to PCO_2_, a response of gap junction channels to PCO_2_, has only been seen for Cx26 ^18^. Surprisingly these GJCs close in response to elevated levels of PCO_2_ rather than opening. Structurally, connexins have a similar architecture with each subunit of the hexameric connexon being built of four transmembrane helices with a cytoplasmic N-terminal helix, involved in gating ^19–21^, pointing into a central pore ^22–28^. The major variations between the sequences of different connexins lie in the length of the cytoplasmic C-terminus, which is denoted in the nomenclature of the connexins, but also in a cytoplasmic loop located between TM2 and TM3 ^29^. In structures of GJCs, the structure on the extracellular side, which is involved in docking between opposing connexons, is well-resolved ^22,25,27,28,30–32^ but is progressively less defined on the cytoplasmic side with neither the C-terminus, nor the cytoplasmic loop being well resolved in any structure. The connexins for which the hemichannels are sensitive to PCO_2_ have a common motif on the cytoplasmic side of the protein that has been shown to be necessary for sensitivity ^13,15,17^ (KΦ(R/K/H)ΦXG) (where Φ corresponds to a hydrophobic residue and X to any residue). Two major conformations of the protein have been elucidated; a pore-constricted state (PCS) in which the N-termini point radially into the pore to form a narrow constriction and a more open state (POS) where the N-terminal helices are flexible ^30,31^. Elevated levels of PCO_2_ bias the conformation of human Cx26 towards the pore constricted conformation ^31^. In the structure of *Gallus gallus* Cx26 (*gg*Cx26) gap junction channel, only the more open form is observed corroborating the functional experiments, which show that *gg*Cx26 gap junction channels do not close in response to PCO_2_ ^33^.

As seen for human Cx26 (hCx26), gap junction channels from *Lepidosiren paradoxa* (*lp*Cx26) are sensitive to PCO_2_ ^34^. However, rather surprisingly, the corresponding hemichannels are not. Relative to other connexins hCx26 has a short cytoplasmic tail. In *lp*Cx26, which has 73% identity to hCx26, the tail is 29 residues longer (Supplementary Figure 1). Its presence has been shown to influence the CO_2_ response of the hemichannels as the replacement of two glycine residues with prolines restores CO_2_ sensitivity ^34^. Similarly the extended C-termini of *Xenopus* and *Latimeria* Cx26 also interfere with hemichannel opening to CO_2_ ^34^. Here we solve the structure of the *lp*Cx26 gap junction channel under the same high PCO_2_ conditions that were used for hCx26. With respect to the other structures of Cx26, TM3 is longer and appears to be stabilised by a detergent molecule, used during the purification. This detergent molecule located between TM3 and TM4 would prevent the conformational changes necessary for the protein to adopt the pore constricted form. In line with this the protein adopts a conformation similar to the open form with the N-terminal helix pulled slightly out of the pore.

## Results

*lp*Cx26 was solubilised and purified in DDM and cryo-EM grids were prepared as for the high PCO_2_ form of hCx26 ^31^. Processing of the data resulted in maps at nominal resolutions of 2.89Å and 2.32Å for C1 and D6 symmetry respectively as defined by gold standard Fourier Shell Correlations (*lp*Cx26-DDM, Supplementary Figure 2-3, Table 1) ^35,36^. Qualitatively the two maps looked similar and as for other Cx26 structures ^9,25,30,31,33,37^, were of much higher resolution in the region corresponding to the extracellular side relative to the cytoplasmic side (Supplementary Figure 2-3). With respect to the published cryo-EM structures of Cx26, the most noticeable feature was additional density as a result of TM3 extending further towards the cytoplasmic side (Figure 1a). While TM4 is also slightly longer than in the other Cx26 structures, there is little density associated with the additional residues of the cytoplasmic tail.

**Figure 1:**
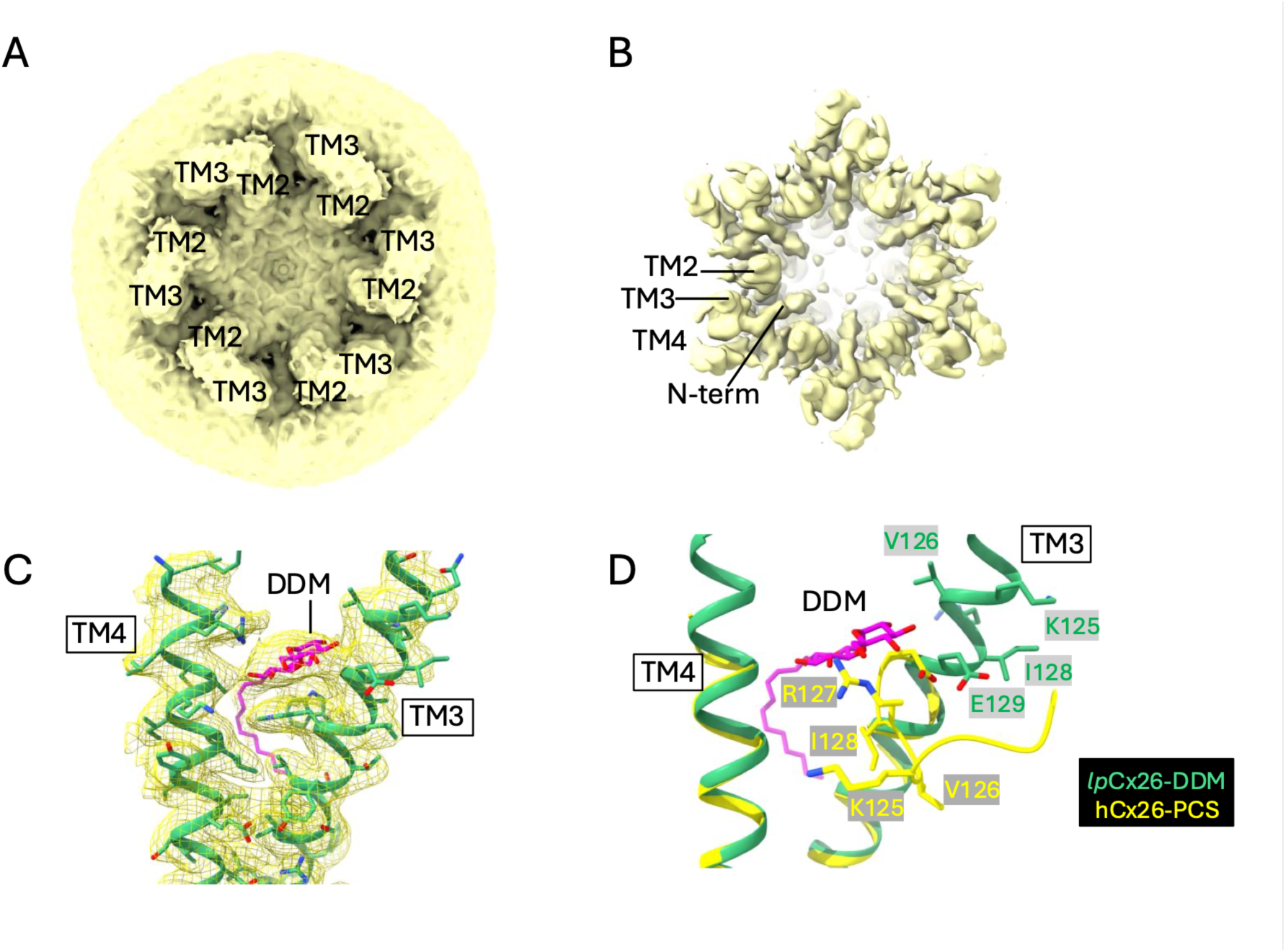
*lp*Cx26 purified in DDM. **A)** Density map showing clear bumps associated with TM2 and TM3 on the cytoplasmic side of the protein. The map has been calculated following refinement with the high resolution information downweighted by using a blurring factor in Coot ^50^. **B)** Map as in A contoured at a higher contour level. **C)** Density associated with the DDM molecule. **D)** Structure of the pore constricted form (PDB 8QA0; yellow) on that of *lp*Cx26-DDM (green).

**Table 1:**
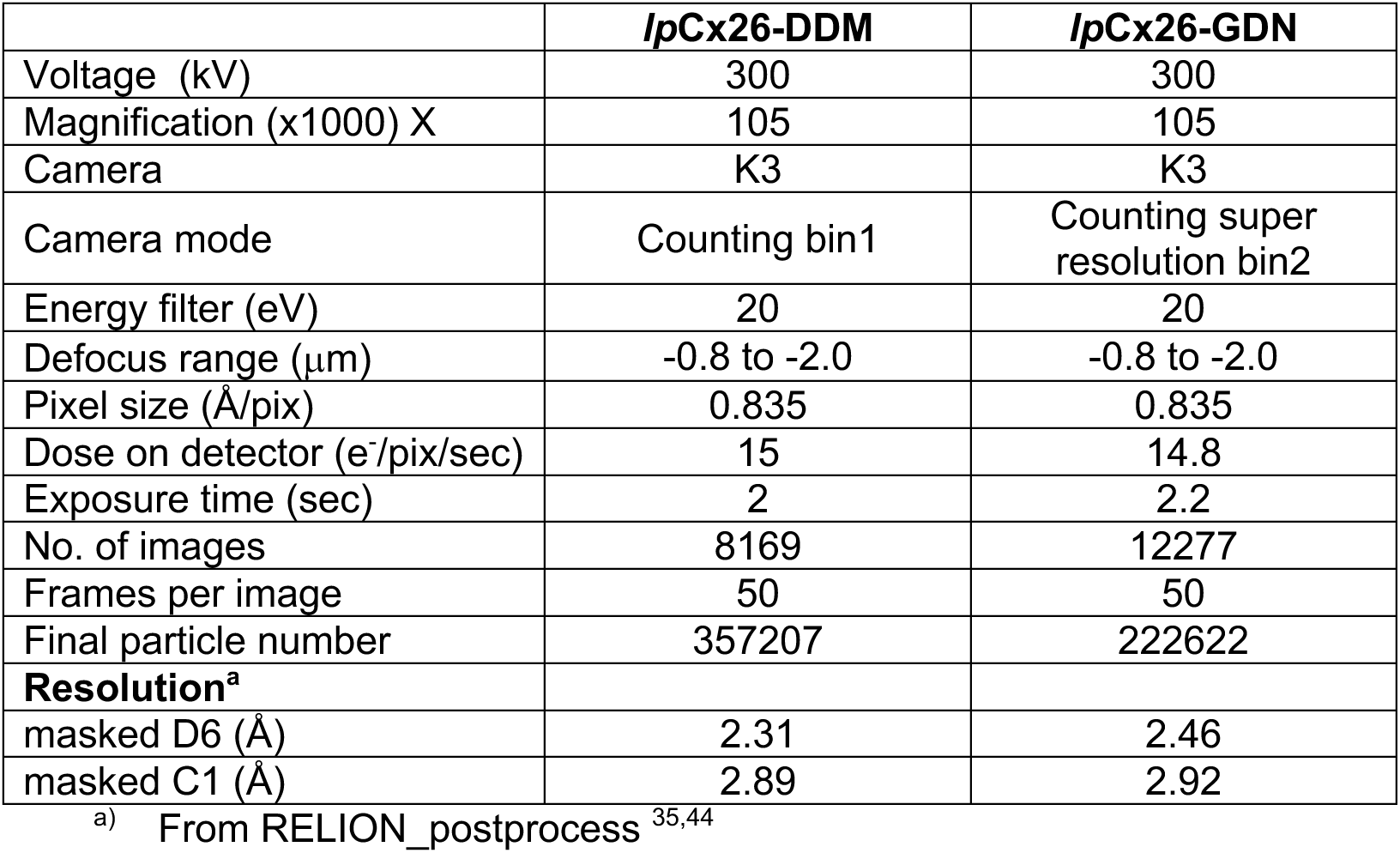
Cryo-EM data collection and processing statistics.

While density was apparent for the N-terminus (Figure 1b), this was less defined than for hCx26 at the equivalent PCO_2_ and was more reminiscent of structures of the gap junction channels under low PCO_2_ conditions ^31^ and from *gg*Cx26 ^33^, where the GJCs do not respond to PCO_2_. More extensive classification of particles based on single subunits or the hemichannel as carried out previously ^30,31^ did not give any significantly different classes. Given that functional assays showed *lp*Cx26 GJCs to be closed at this level of PCO_2_, this was surprising. However, in building the structure we observed density between TM3 and TM4 that was consistent with the sugar moiety of a detergent molecule (n-dodecyl-β-D-maltopyranoside; DDM) (Figure 1c). The molecule in this position would not be compatible with the pore constricted conformation of the protein that we observed for hCx26 (Figure 1d).

We therefore collected a further cryo-EM data set (Supplementary Figure 3-4; Table 1) by replacing DDM with glyco-diosgenin (GDN) during the final size exclusion step. GDN, although having the same sugar backbone would be more conformationally restricted. Refinement resulted in a map at 2.48Å with D6 symmetry imposed that was essentially identical to the map obtained with DDM, with similar density between TM3 and TM4. Further classification focussed on the hemichannel (Supplementary Figure 4; Table 2) resulted in two maps with slightly different appearance that could be refined to 2.71Å and 2.89Å respectively. In one, the N-terminus (*lp*Cx26-GDN-1) was better defined pointing into the pore allowing a tentative placement of the residues (Figure 2a). In the second (*lp*Cx26-GDN-2), the N-terminus was poorer, with some evidence of a second conformation parallel to the membrane, but the density associated with TM3 extended further to the cytoplasmic side with respect to *lp*Cx26-GDN-1 (Figure 2b). However, overall, the conformation of the protein was similar with just subtle differences between them (RMSD across all matching Cα pairs of 0.460Å) so that no significant conformational change could be associated with the resulting maps. Focussing the classification on just one subunit was qualitatively similar. Both *lp*Cx26-GDN-1 and *lp*Cx26-GDN-2 showed similar density for the detergent molecule (Supplementary Figure 5a). While this may have been from GDN, our inference, based on the density (Supplementary Figure 5a) was that this was more likely to be DDM that did not exchange with the GDN. This suggests a relatively tight binding site, with the detergent occupying the binding site before the exchange into the CO_2_/HCO_3_^-^ buffer.

**Figure 2:**
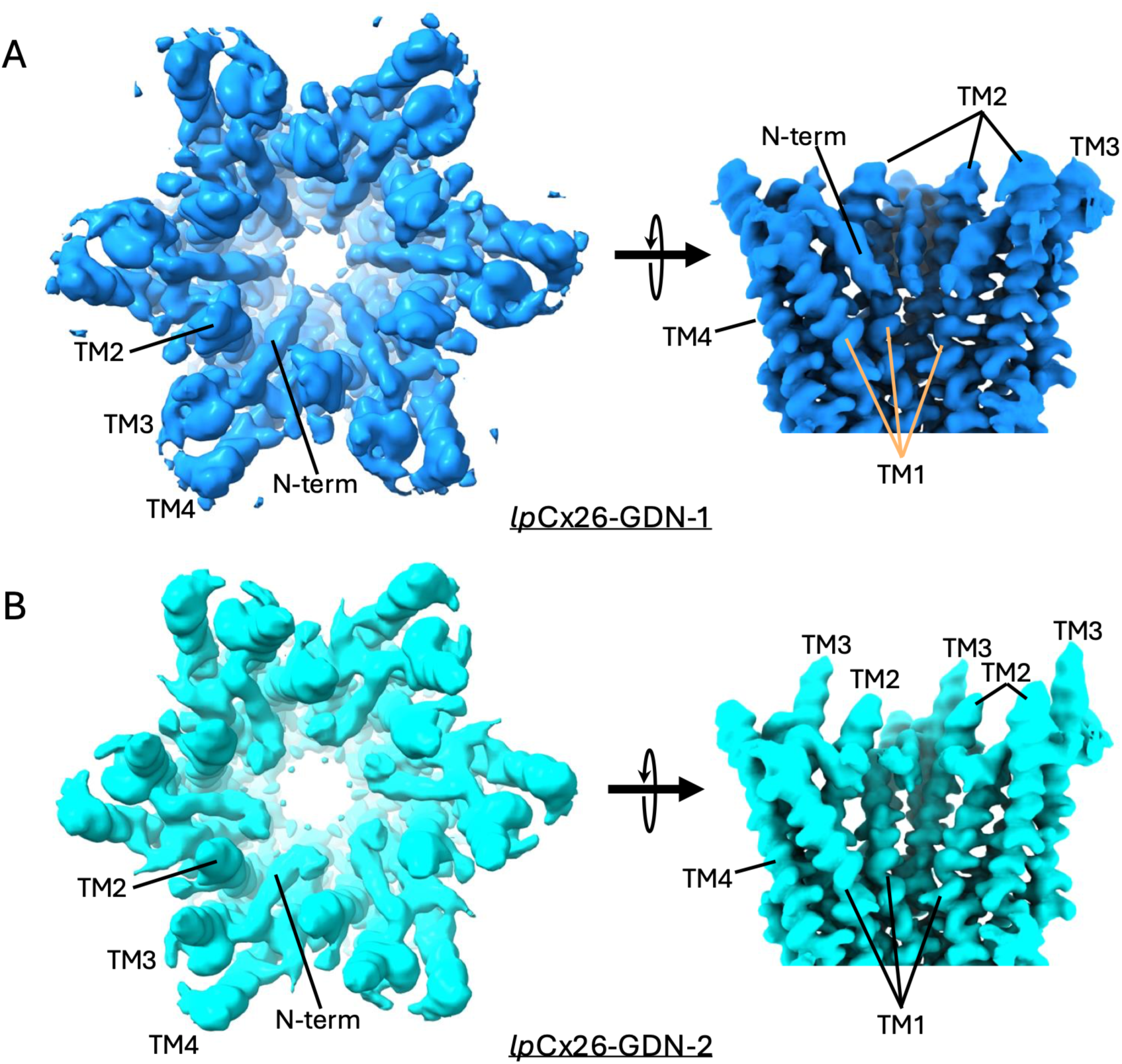
*lp*Cx26 exchanged into GDN. **A**) In *lp*Cx26-GDN-1 the N-terminus is better defined but there is less density for TM3. **B**) In *lp*Cx26-GDN-2 the N-terminus is less clear but TM3 extends further. A blurring factor has been applied to both maps to clarify the overall envelope as in Figure 1.

**Table 2.** Cryo-EM refinement and validation statistics.

|  | *lpCx26-GDN-1* | *lpCx26-GDN-2* |
| --- | --- | --- |
| Deposited Structure PDB ID | **28MC** | **28ME** |
| Final particle number | 57156 | 71467 |
| **Map resolution** |  |  |
| FSC threshold | 0.143 | 0.143 |
| Symmetry imposed | C6 | C6 |
| RELION postprocess <sup>a</sup> | 2.7 | 2.9 |
| Phenix Unmasked (Å) <sup>b</sup> | 2.4 | 2.5 |
| Phenix Masked (Å) <sup>b</sup> | 2.4 | 2.5 |
| **Refinement** |  |  |
| Initial model | Alphafold | *lpCx26-GDN-1* |
| Resolution (Å; FSC=0.5) Model | 2.6 | 2.7 |
| Sharpening B factor (Å<sup>2</sup>) | local | local |
| **Model composition** | 12 chains | 12 chains |
| Non-hydrogen atoms | 21084 | 21456 |
| Protein residues | 2400 | 2472 |
| Water | 288 | 312 |
| Ligand: lipid/detergent | 36 | 36 |
| **B factor (Å<sup>2</sup>)** |  |  |
| protein | 54 | 46 |
| water | 33 | 34 |
| Lipid/detergent | 84 | 61 |
| **R.m.s. deviations** |  |  |
| Bond lengths (Å) | 0.005 | 0.004 |
| Bond angles (°) | 0.992 | 0.936 |
| **Validation** |  |  |
| MolProbity score | 2.04 | 1.94 |
| Clashscore | 7.88 | 11.55 |
| Rotamer outliers (%) | 3.30 | 2.97 |
| **Ramachandran plot** |  |  |
| Favoured (%) | 97.19 | 99.01 |
| Allowed (%) | 2.81 | 0.99 |
| Disallowed (%) | 0 | 0 |
| **CaBlam Outliers (%)** | 1.56 | 1.01 |
| **Correlation coefficients** |  |  |
| CC (mask) | 0.88 | 0.87 |
| CC (box) | 0.71 | 0.71 |
| CC (peaks) | 0.69 | 0.67 |
| CC (volume) | 0.86 | 0.83 |
| Mean CC for ligands | 0.72 | 0.71 |

a) From postprocess job in RELION 5
b) From Phenix Validation job <sup>48</sup> using locally sharpened maps <sup>47</sup>

With respect to the constricted form of hCx26 ^30^ the N-terminal helices are pulled back from the centre of the pore (Figure 3a-c). While the N-terminal residues could only be tentatively assigned in the density, the loop between TM1 and the N-terminus is very clear with Val13 packing onto Ala92 (Supplementary Figure 5b). Overall, the conformations of the C-terminal residues of TM1 and preceding loop are most similar to the structure of *gg*Cx26 ^33^ rather than the rotated conformation seen in the pore constricted form (Figure 3a). This position of the loop would not be compatible with the position of TM2 in the hCx26 pore-constricted form (Figure 3b).

**Figure 3:**
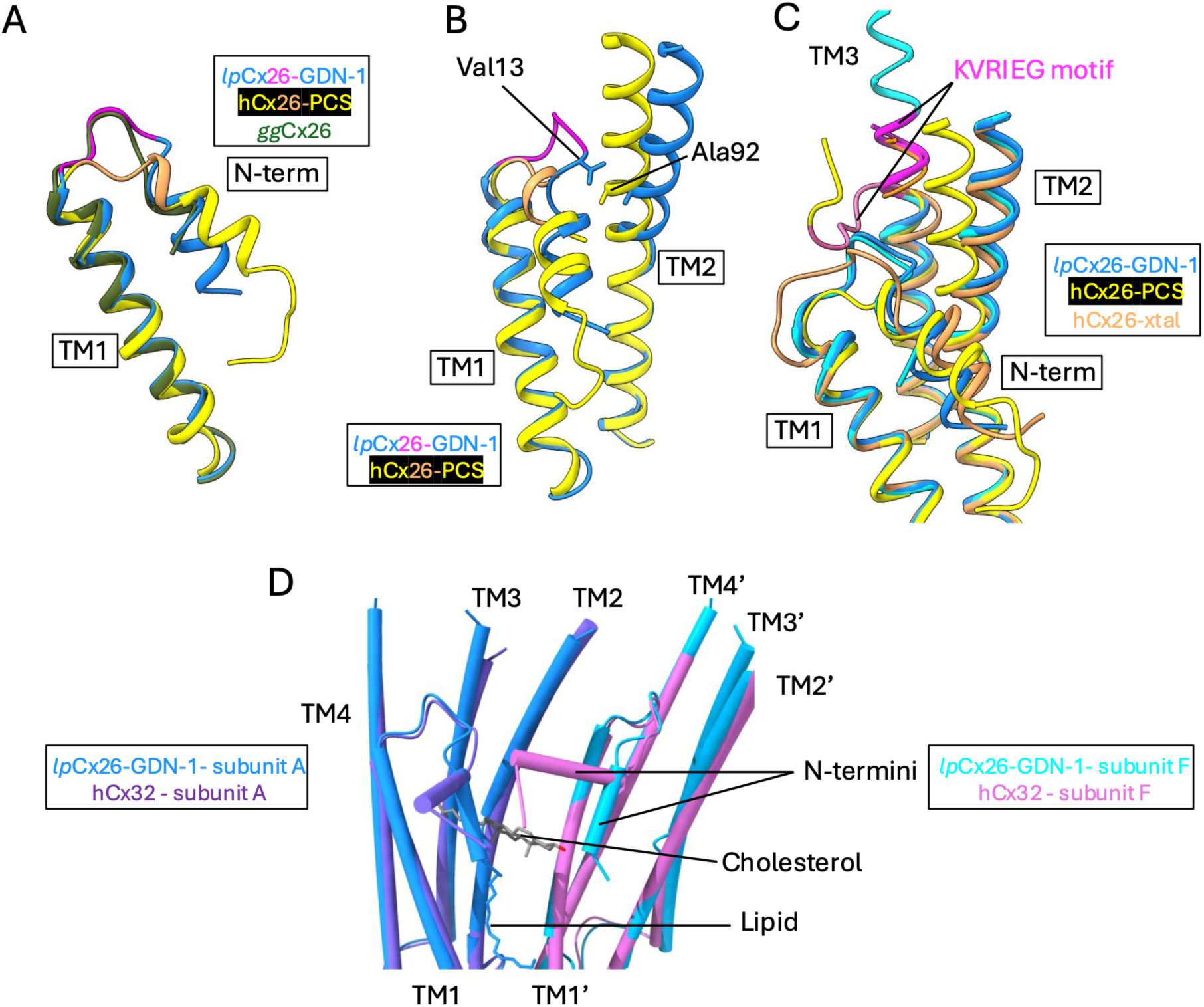
Comparison of *lp*Cx26 with other structures of Cx26 and Cx32. **A)** Comparison of the N-terminus and TM1. *lp*Cx26-GDN-1 is shown in dodger blue, the pore constricted form of hCx26 (PDB 8QA0) in yellow and *gg*Cx26 (8QVN) in olive. The loop joining the N-terminal helix and TM1 is shown in magenta for *lp*Cx26 and goldenrod for hCx26. **B)** Similar to A but highlighting the potential clash of Val13 with Ala92 were TM2 of *lp*Cx26 to adopt the same conformation as seen in hCx26 without any adjustments of the N-terminus. **C)** Comparison of the two lpCx26 structures (dodger blue and cyan), the pore constricted form of hCx26 (8QA0; yellow) and the crystal structure (2ZW3; goldenrod). The residues of the KVRIEG motif are shown in magenta for *lp*Cx26 and hot pink for hCx26. The superposition of the structures was based on the A molecule. **D)** Comparison of *lp*Cx26-GDN-1 with hCx32 in lipid nanodiscs with cholesterol hemisuccinate (PDB 9QNF). Two neighbouring molecules of the dodecamer are shown coloured dodger blue and cyan for *lp*Cx26-GDN-1 and purple and pink for Cx32. The modelled cholesterol is shown for hCx32, binding under the N-terminal helices. The lipid that is seen in Cx26 and other connexins is shown in dodger blue. The lipids associated with Cx32 that fill the pore have been omitted for clarity.

TM3 is much longer than seen in the other cryo-EM Cx26 structures so that the helix has been modelled from Ser117 in *lp*Cx26-GDN-2 (Figure 3c) and from Val126 in *lp*Cx26-GDN-1. In both hCx26 and *gg*Cx26 cryo-EM structures, the first residue assigned to TM3 is Gly130, which is the final residue in the KVRIEG motif important in sensitivity to CO_2_. In the pore-open structure, the residues of the KVRIEG motif are disordered, whereas in the pore-constricted conformation they form a linear peptide. That these residues form part of TM3 is more similar to the crystal structure of hCx26 ^25^. In this latter structure the overall conformation of the protein is, in fact, most similar to the open conformation that we observe in *lp*Cx26, based on the position of TM1, TM2 and the N-terminus (Figure 3c).

The hydrocarbon chains of other detergent or lipid molecules are also evident in the density. Two of these have been modelled in the final structure (Supplementary Figure 6a). As seen for hCx26 and *gg*Cx26 ^30,31,33^, density that would be consistent with a lipid molecule is present in all maps within the pore of the protein (Supplementary Figure 6). Density is also clear for a lipid binding between TM1 and TM4 of one subunit and TM2 and TM3 of the next (Supplementary 6). Given that the headgroup is reasonably clear this has been modelled as phosphatidylethanolamine. In this position TM3 is sandwiched between the lipid headgroup and the DDM and the lipid tails surround the side-chain of Trp24.

Recently structures have been reported for human Cx32 gap junction channels (hCx32) ^22,38^. Cx32, like Cx26, is a β connexin. It shares 63% identity within the core region compared to both hCx26 and *lp*Cx26 (Supplementary Figure 1). When the structure was solved from protein in detergent the GJC has a more disordered N-terminus as we have seen for the low PCO_2_ hCx26 and *gg*Cx26 and TM3 only starts at Gly129 ^22^. When solved from protein in lipid nanodiscs and cholesterol hemisuccinate the density on the cytoplasmic side is much better resolved ^38^. Although this is not modelled in the associated structure, in the deposited map (EMD-53245) TM3 can be built with a high level of confidence from residue 120 (Supplementary Figure 7). Density between TM3 and TM4 also indicates the presence of a small molecule in the same position as the DDM that we see in *lp*Cx26 (Supplementary Figure 7). Overall, the conformation of the lpCx26 is very similar to that of *hCx32* with an rmsd of 0.9Å over all 189 matched pairs common to the structures (Figure 3d). The major difference between the structures is in the position of the N-terminus. In the Cx32 structure in nanodiscs with cholesterol hemisuccinate bound, the N-terminal helix lies parallel to the membrane pointing to the side of the pore in an anti-clockwise fashion looking from the cytoplasmic side. Instrumental in stabilising this conformation is cholesterol hemisuccinate which binds below the helix ^38^. The loop joining the N-terminal helix to TM1, however is very similar between *lp*Cx26 and Cx32 (Figure 3d).

## Discussion

We have previously elucidated two major conformations of human Cx26: a conformation in which the pore is constricted by the N-terminus and one in which the N-terminus is more flexible so that the aperture to the pore is more open ^30,31,33^. The occurrence of these states correlates with the functional studies ^18^ in that at high PCO_2_ where the gap junction channels are closed the equilibrium is pushed towards the constricted state and at low PCO_2_ or where the channels are not responsive to CO_2_ the more open conformation is observed. Here we observe an apparent discrepancy in that functional studies show that *lp*Cx26 gap junctions close in response to high PCO_2_ ^34^ and yet a more open conformation of the protein is observed. With respect to previous cryo-EM structures of Cx26 in the open form the N-terminus is slightly more defined but, though it still points into the pore, is pulled back from the centre relative to the pore-constricted conformation. The discrepancy between the structure and functional studies may be due the binding of a putative detergent molecule between TM3 and TM4. In this position, the detergent molecule would prevent the conformation seen in the constricted form of the protein by sterically hindering residues 122 to 130 including the KVRIEG motif from adopting the linear conformation that is seen in the constricted form. Thus, the DDM molecule may be trapping the protein in the more open state. Given that the density looked similar even after replacement with GDN, it seems that the DDM binds tightly during solublisation and does not exchange during size exclusion. A similar situation may have occurred in the recently reported structures of hCx32 ^38^ where we observe density in the same position in the associated maps, which could be modelled as DDM. As for *lp*Cx26, DDM was used at high concentrations during solubilisation of the Cx32 ^38^ but was not present in the final purification steps. In human Cx26 a detergent binding in this position would be sterically hindered by the side-chain of Tyr217. In both lpCx26 and hCx32 the tyrosine is replaced with an alanine. Overall, the conformation of the hCx32 structure is very similar to that of *lp*Cx26, the main difference being in the position of the N-terminal helix that runs parallel to the membrane in Cx32. Its position in Cx32 is influenced by the presence of lipids and cholesterol hemisuccinate ^38^. Given the similarity of the loop following the N-terminus, it is possible that Cx26 could also adopt this conformation of the N-terminus, though, as can be seen from the sequence (Supplementary Figure 1), the residues of the N-terminus differ slightly between the proteins. In particular, in the Cx32 structure, the hydroxyl oxygen of Ser 11 forms hydrogen bonding interactions to stabilise the change in direction of the main chain and Tyr 7 packs between Ser 11 and Ala 96 forming a hydrogen bond with His100. In the Cx26 sequences Ser11 has been replaced with a glycine and Tyr7 with an asparagine. The conformation of the N-terminus in Cx32 is also stabilised by the lipid and cholesterol hemisuccinate in the structure. It would be possible for cholesterol to bind in the same position in Cx26, though the arrangement or type of lipids would need to differ slightly due to the alterations in the side chains of the N-terminus.

The response of hCx26 to CO_2_ has been attributed to carbamylation of Lys125 ^13,18^ of the KVRIEG motif. This labile and consequently reversible covalent modification causes a neutral lysine to become negatively charged by the addition of CO_2_ ^39^. Mutation of Lys125 to alanine prevents the opening of hemichannels ^13^. Mutating the same residue to arginine, which would have the same charge as a protonated lysine prevents the closure of gap junction channels in response to PCO_2_, though it still closes upon acidification ^18^. Conversely, mutating the same residue to glutamic acid, and so mimicking the charge of a carbamylated lysine, causes hemichannels to be constitutively open and gap junctions to be constitutively closed ^13,30^. Whereas, in the pore-constricted form of the protein these residues form a linear peptide, in the *lp*Cx26 structure, as seen in the crystal structure of hCx26 ^25^ and the related structure of hCx32 ^38^, the same residues form part of TM3. The hypothesis was that upon carbamylation, the now negatively charged lysine would form a salt bridge with Arg104 of TM2 and in so-doing cause a conformational change that would close the pore ^13^. Extensive mutational studies back up this hypothesis ^13,18,30,40^. In particular, in studies of hemichannels, replacing both Lys125 and Arg104 with cysteines causes a response to redox potential that can be interpreted as the formation of a disulphide bond between the two residues ^40^ mimicking the response of the wild-type protein to CO_2_. Arg104 is at the cytoplasmic end of TM2. Our structural studies of the gap junction channels show that TM2 undergoes a large conformational change between the pore-constricted and open conformations changing the position of Arg104^30^. It would seem from the *lp*Cx26 structure, backed up by the crystal structure of hCx26 and cryo-EM structures of Cx32 that Lys125 forms part of TM3 in the open conformation. It is interesting to note though, that through classification of hCx26 based on a single subunit a conformation was observed where both the residues preceding the KVRIEG motif as well as those following were helical so that the KVRIEG motif effectively forms a break in TM3 ^30^ (Supplementary Figure 8). Although this is modelled with low confidence, a similar situation is also observed in Alphafold3 ^41^ predictions of *lp*Cx26 (Supplementary Figure 8). Given that in most connexin gap junction channel structures including Cx43 ^27,42^, Cx46/50 ^28,32^ and Cx36 ^23^ TM3 extends only approximately as far as the equivalent of Gly130, which is conserved amongst the CO_2_ sensitive connexins, this region is obviously very flexible. Carbamylation of Lys125 may either cause the transition or trap the constricted conformation. While given the positions of Lys125 and Arg104, the salt-bridge hypothesis is still plausible other residues may be necessary. We have observed, for example, that the mutation of Lys108 to arginine in hCx26 also prevents gap junction closure, though interestingly has no effect on the hemichannel ^33^.

As for hCx26 and *gg*Cx26 we observe a lipid molecule binding in the centre of the pore in this structure of lpCx26. A molecule at this position is present in all the conformations of the protein that we have elucidated ^30,31,33^ and is also present in other cryo-EM structures ^22–24,27,42^. Recent reports of Cx32 ^38^ and other connexins ^23,27,42^ of the conformation of the N-terminus being influenced by the lipid environment have suggested that the movement of lipids and/or cholesterol into and out of the pore could be important in the closure mechanism. We cannot rule out such a mechanism for Cx26. The lipid that we observe within the pore does not change position regardless of the conformation of the protein. The lipid that we have modelled on the outer surface of the pore is also observed in the other Cx26 structures that have a similar conformation, though is not as well defined. On the other hand, the movement of TM2 and Trp24 that occurs in forming the pore constricted structure leads to displacement of the lipid ^30^. Thus, the presence of this lipid may stabilise one conformation of the protein. Whether or not lipids are involved in pore closure, a trigger is still needed to allow molecules to move into or be removed from the pore aperture. The position of TM1 and TM2 in the pore-constricted state would not be compatible with the position of cholesterol seen in hCx32 so that the cholesterol molecules would necessarily have to move their positions.

With the elucidation of this structure, we are now able to refine our previously reported mechanism (Figure 4). When the KVRIEG motif forms part of TM3 the N-terminal helix is pulled further out of the pore opening the aperture. Concerted movements in TM3, TM2 and TM1 serve to push the N-terminal helix further into the pore. The presence of the detergent molecule located between TM3 and TM4 appears to stabilise TM3 and prevent the conformational changes that would allow pore constriction. It could easily be envisaged that slight modifications to the cytoplasmic tail could have the same effect. Blocking of conformational changes with small molecules in this way could provide a means of modulating connexins.

**Figure 4:**
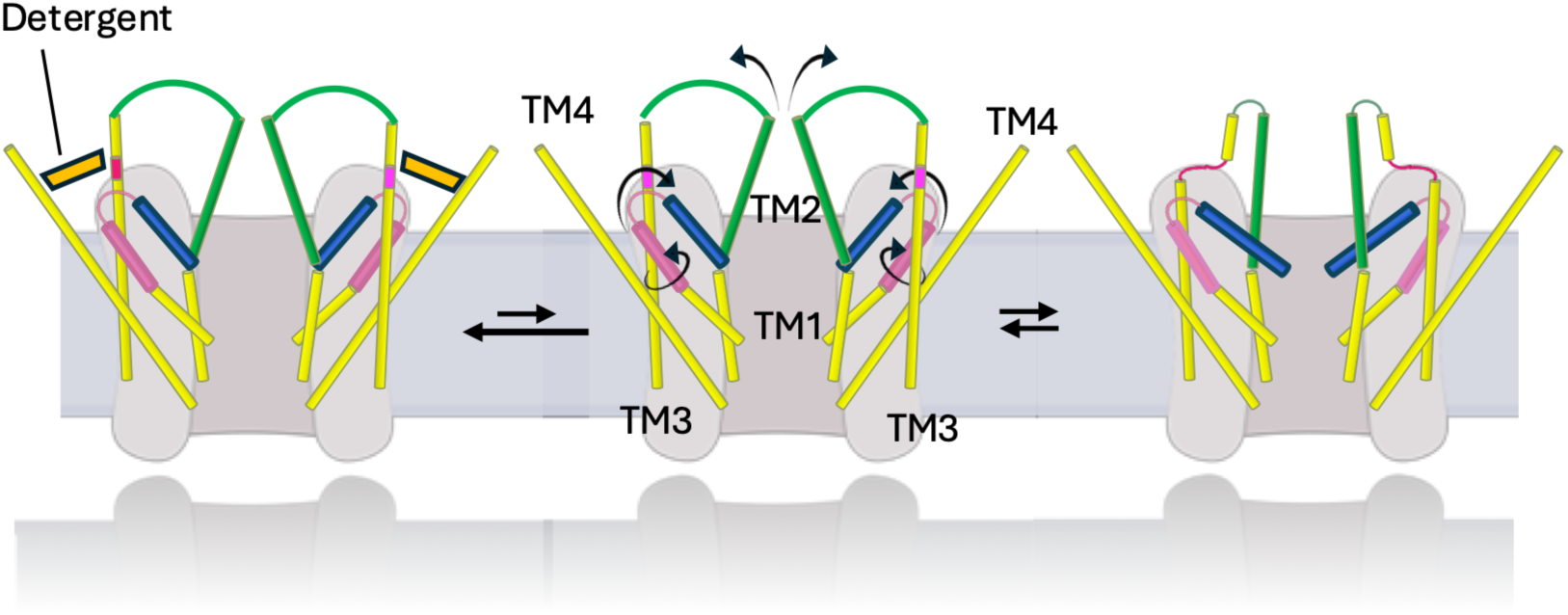
Schematic of mechanism. Cx26 exists in an equilibrium between a pore open form (middle) and a pore-constricted form (right). In changing conformation, the cytoplasmic part of TM2 (green) flexes and that of TM1 (pink) rotates necessitating changes to the N-terminus (blue) ^30^. Associated with these changes is the conformation of the KVRIEG motif (magenta). In the pore-constricted form these residues form a linear peptide, whereas the same residues can form part of TM3 when the N-terminus is pulled back. Detergent (orange quadrangle) binding between TM3 and TM4 appears to stabilise the open conformation (left).

## Methods

### Cloning and expression of lpCx2C

DNA encoding for *lp*Cx26 lacking a stop codon was prepared by PCR from a *lp*Cx26-mCherry expression construct described previously ^34^, using primers which introduced a C-terminal His-6 purification tag. (forward primer: 5’-CACGACGAATTCCACCATGAACTGGGGTAC-3’, reverse primer 5’-GCTATCACTAGTAGCTTGGTTGGTAAGG-3’).

The amplified lpCx26 sequence was double digested with EcoRI and SpeI, prior to ligation into the corresponding sites of the pFastbac1 vector (NEB). The construct was sequence verified before being transformed into DH10Bac cells (Invitrogen) for bacmid production, and subsequent baculovirus production in Sf9 cells, according to the manufacturer’s instructions. Sf9 cells were harvested 72 hours post infection for protein purification.

### Initial Purification

The purification was performed as for hCx26 described previously ^43^, with the following changes: cells were broken with 20x strokes with a dounce homogeniser, solubilisation was for 3.5 hours, and dialysis was carried out against (20 mM sodium phosphate, 500 mM NaCl, 5 % glycerol, 1 mM DTT, 0.03% DDM (Glycon Biochemicals GMBH), pH 7.4) to remove histidine, before concentrating the protein to ∼3mg/ml and flash-freezing the protein.

### *lp*Cx26-DDM

#### Cryo-EM sample preparation and data collection

The flash-frozen protein was thawed centrifuged at 21,000 x g 4°C, 30mins prior to concentrating in a viva6 100kDa MWCO to 18mg/ml for size exclusion chromatography using a Superose 6 5/150 column (GE Healthcare Lifescience) to exchange the buffer to *S0 mmHg aCSF buffer*: (70 mM NaCl, 5 % glycerol, 1 mM DTT, 0.03 % DDM, 80 mM NaHCO_3_, 1.25 mM NaH_2_PO_4_, 3 mM KCl, 1 mM MgSO_4_, 4 mM MgCl_2_.) The *lp*Cx26 peak was concentrated to 4.3 mg/ml before being gassed with the correct amount of 15% CO_2_ to give a final pH of ∼7.4 as previously described ^43^. Aultrafoil 300 mesh gold grids 0.6/1 (Quantifoil Micro Tools GMBH) were glow discharged 35 mA, for 60 seconds prior to use. Vitrification of the protein in liquid ethane at −180°C was carried out with a FEI Vitrobot with 3 μl protein per grid at 4°C, 100 % humidity, 3 seconds at blotforce 10 in a 15% CO_2_ / N_2_ atmosphere. Data were collected on an FEI Titan Krios G3 equipped with a K3 detector and BioQuantum energy filter using a 20 eV slit width. A dose rate of ∼15 e/pix/sec on the detector was chosen with a final dose of 43 e/Å^2^ on the specimen. Data collections were prepared and run automatically using EPU3.0 and aberration-free image shift (AFIS).

#### Cryo-EM data processing

Data were processed using RELION4 ^44^. Micrographs were motion corrected using the version of MotionCor2 ^45^ implemented in RELION, and CTFs were estimated using CTFFIND4 ^46^. Particles were picked using the Laplacian of Gaussian (LoG) picker, and after 3 rounds of 2D classifications with particles downsampled to 4Å/pixel, good particles were selected. 3D classification was carried out in C1 with an initial model generated from hCx26 K125R ^30^ with a low-pass filter of 40Å. 3D classification in C1 resulted in 3 similar classes, which were selected for continued processing including further 2D classification. Particles were then unbinned and subjected to refinement, CTF refinement and polishing in RELION4 to improve the resolution of the Coulomb shells until no further improvement was gained. The resolution was estimated based on the gold standard Fourier Shell Coefficient (FSC) criterion ^35,36^ with a soft solvent mask. Local Resolution estimation was carried out in RELION4. The map was sharpened using local sharpening ^47^ in Phenix ^48^. Attempts at subclassifying the particles further were made following previous protocols for focussed classifications ^30^ based on either the cytoplasmic regions of the hemichannels or of single subunits. An initial model was created in Alphafold2 ^49^ and built into the density using Coot ^50^.

### *lp*Cx26-GDN

#### Cryo-EM sample preparation and data collection

The flash-frozen sample was defrosted, centrifuged at 21,000xg 4°C, 30mins prior to concentrating in a viva6 100kDa MWCO to 18mg/ml. The sample was re-centrifuged as above prior to injection onto Superose6 5/150 equilibrated with *S0 mmHg aCSF buffer*: (70 mM NaCl, 1 mM DTT, 0.006 % GDN, 80 mM NaHCO_3_, 1.25 mM NaH_2_PO_4_, 3 mM KCl, 1 mM MgSO_4_, 4 mM MgCl_2_.)

The lpCx26 peak was taken directly off the column at a concentration of 1.6 mg/ml, and gassed as above. Aultrafoil 300 mesh gold grids 1.2/1.3 (Quantifoil Micro Tools GMBH) were glow discharged 35 mA, for 180 seconds prior to use. Vitrification of the protein in liquid ethane at −180°C was carried out with a FEI Vitrobot with 3 μl protein per grid at 4°C, 100 % humidity, 3 seconds at blotforce 10. Data were collected on an FEI Titan Krios G3 equipped with a K3 detector and BioQuantum energy filter using a 20 eV slit width. A dose rate of ∼14.8 e/pix/sec on the detector was chosen with a final dose of 47 e/Å^2^ on the specimen. Data collections were prepared and run automatically using EPU3.8 or 3.9 and aberration-free image shift (AFIS).

#### Cryo-EM data processing

Data were processed using RELION5. Micrographs were motion corrected using the version of MotionCor2 ^45^ implemented in RELION5, and CTFs were estimated using CTFFIND4 ^46^. Particles were picked using reference-based template matching in RELION5 based on references from 2D class averages produced from a total of 950 particles that were manually picked from 66 micrographs. After 1 round of 2D classifications with particles downsampled to 4Å/pixel, good particles were selected. An *Ab initio* model was created and a 3D classification was carried out in C1 with the initial model low-passed filtered to 30Å. A further round of 3D classification including Blush regularisation ^51^ resulted in a single class. Particles were then unbinned and subjected to refinement with Blush regularisation, CTF refinement and polishing in RELION5. The resolution was estimated based on the gold standard Fourier Shell Coefficient (FSC) criterion ^35,36^ with a soft solvent mask. Focussed classification was carried out based both on a single subunit and for the hemichannel as described previously ^30,31^. Only the results from the hemichannel classification were taken further. For this classification, following particle expansion, 3D classification with Blush regularisation and C6 symmetry imposed was carried out with a mask focussed on the cytoplasmic region of the hemichannel. Two classes were selected for further analysis based on the clarity of the features. The particles were unsubtracted, duplicates removed and unmasked local refinement with Blush regularisation was carried out with C6 symmetry imposed. Local Resolution estimation was carried out in RELION5. The maps were sharpened using local sharpening^47^ in Phenix ^48^.

#### Model building

The preliminary model built into the *lp*Cx26-DDM map was used as the initial model for refinement against the pCx26-GDN-1 map. Model building was carried out in Coot ^50^, with real space refinement in Phenix ^48^ against the sharpened maps using NCS constraints. Density on the outer face of the pore was tentatively modelled as phosphotidylethanolamine and DDM was inserted into density located between TM3 and TM4. Density within the pore was modelled with a short hydrocarbon chain. Water molecules were inserted manually in Coot.

## Structural Analysis

Structural images shown in this paper were generated in ChimeraX ^52,53^. Superpositions were carried out in ChimeraX such that only matching Cα pairs within 2Å after superposition were included in the matrix calculation. AlphaFold3 models shown in Supplementary Figure 8 were created with AlphaFold3 ^41^ on the AlphaFold server.

## Acknowledgements

We thank the Leverhulme Trust (RPG-2015-090) and MRC (MR/P010393/1) for support. We acknowledge the Midlands Regional Cryo-EM Facility, hosted at the Warwick Advanced Bioimaging Research Technology Platform, for use of the JEOL 2100Plus, and the Midlands Regional Cryo-EM Facility, hosted at Leicester Institute of Structural and Chemical Biology for use of the FEI Titan Krios G3 and associated facilities, supported by MRC award reference MC_PC_17136. We thank Dr TJ Ragan, Dr C. Savva and Dr E. Hesketh for help with cryo-EM. We are grateful to the technical support in the School of Life Sciences, University of Warwick.

## Author Contributions

The project was initiated and supervised by ND and AC. Cloning, expression, purification and grid preparation were carried out by DB. Data collection was performed by DB. Data processing was done by DB and AC. DB and AC refined and analysed the structures. AC, wrote the manuscript with help from all authors.

## Data Availability

Cryo-EM density maps have been deposited in the Electron Microscopy Data Bank (EMDB) under accession numbers EMD-56611 (*lp*Cx26-DDM), EMD-56608 (*lp*Cx26-GDN-1), EMD-56609 (*lp*Cx26-GDN-2). Structure models have been deposited in the RCSB Protein Data Bank under accession numbers 28MC (*lp*Cx26-GDN-1) and 28ME (*lp*Cx26-GDN-2).

## Supplementary Figures

**Supplementary Figure 1:**
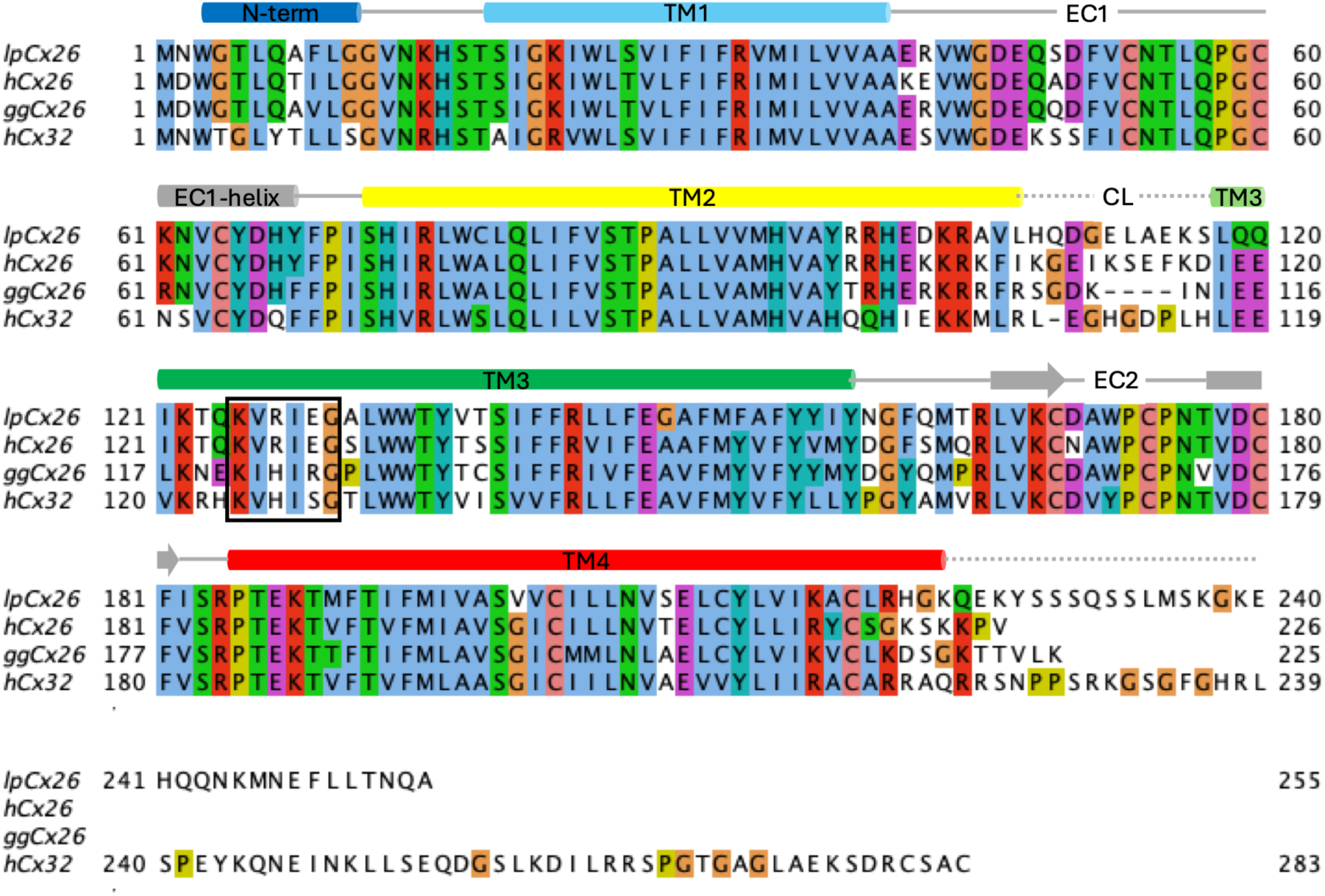
Sequence comparison of *lp*Cx26 (GEHZ01053112.1), hCx26 (NP_003GG5.2), *gg*Cx26 (NP_001257745.1) and hCx32 (NP_000157). The sequence has been coloured according to the Clustal palette in Jalview ^54^. The secondary structure is based on the structure of lpCx26 and the helices are coloured from blue to red based on the rainbow algorithm implemented in ChimeraX ^53^. The residues of the KVRIEG motif are boxed. *lp*Cx26 shares 74% sequence identity with hCx26, 70% with *gg*Cx26 and 63% with hCx32.

**Supplementary Figure 2:**
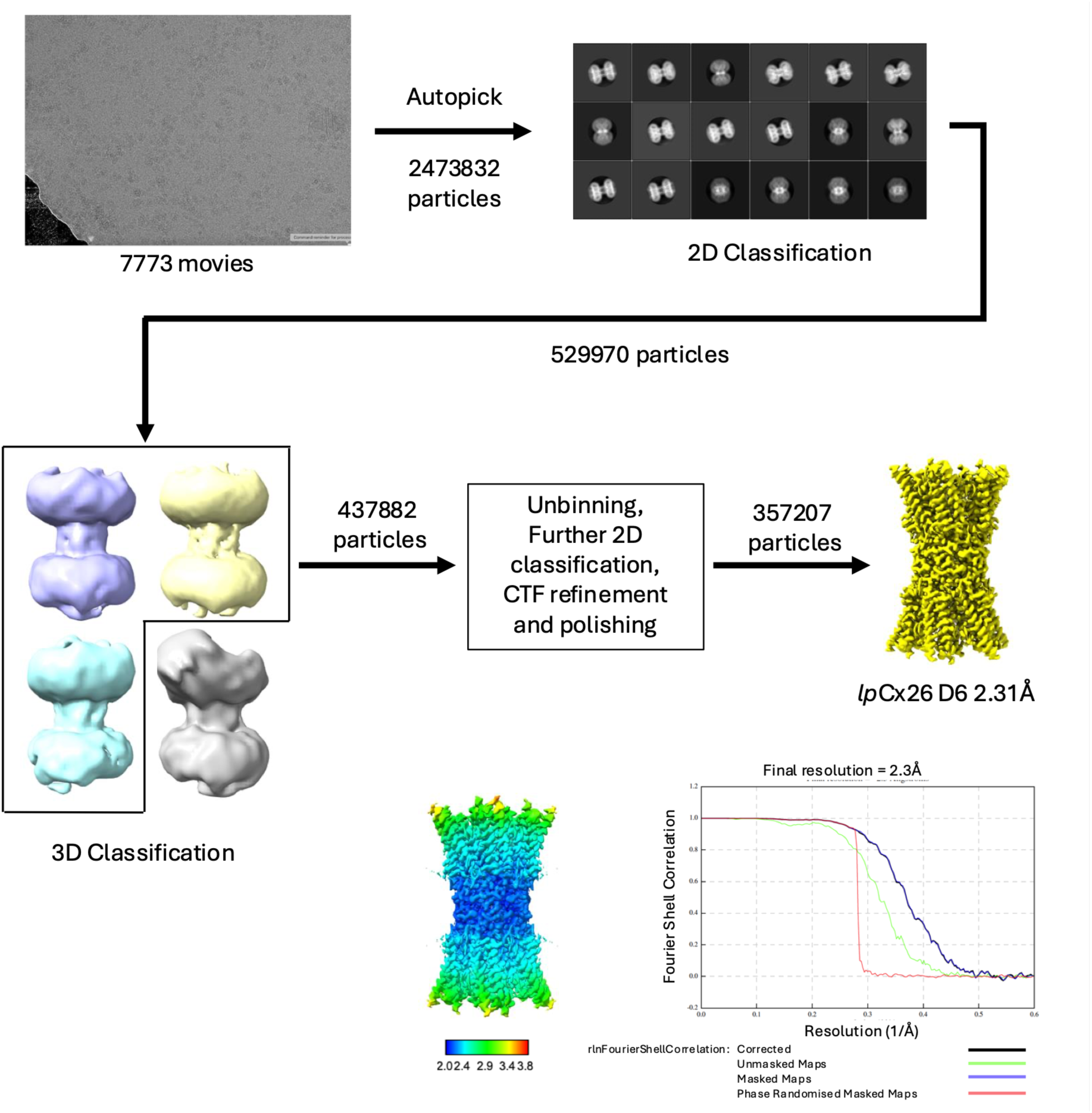
Workflow for processing of cryo-EM data for *lp*Cx26 purified in DDM.

**Supplementary Figure 3:**
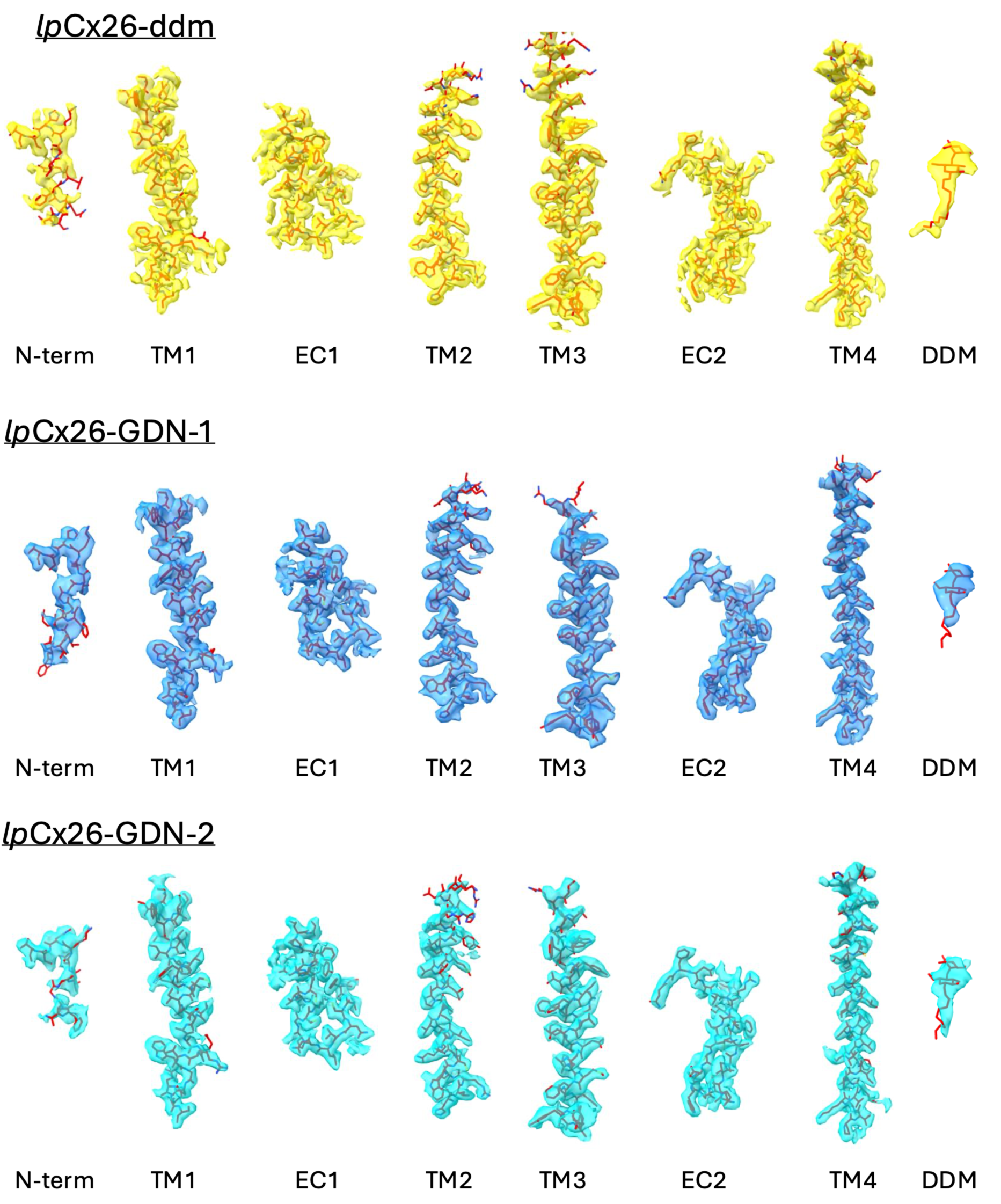
Density associated with the structures. Density corresponding to the N-terminal region (N-term), transmembrane helices (TM1-4), extracellular loops (EC1, EC2) and the DDM for each structure.

**Supplementary Figure 4:**
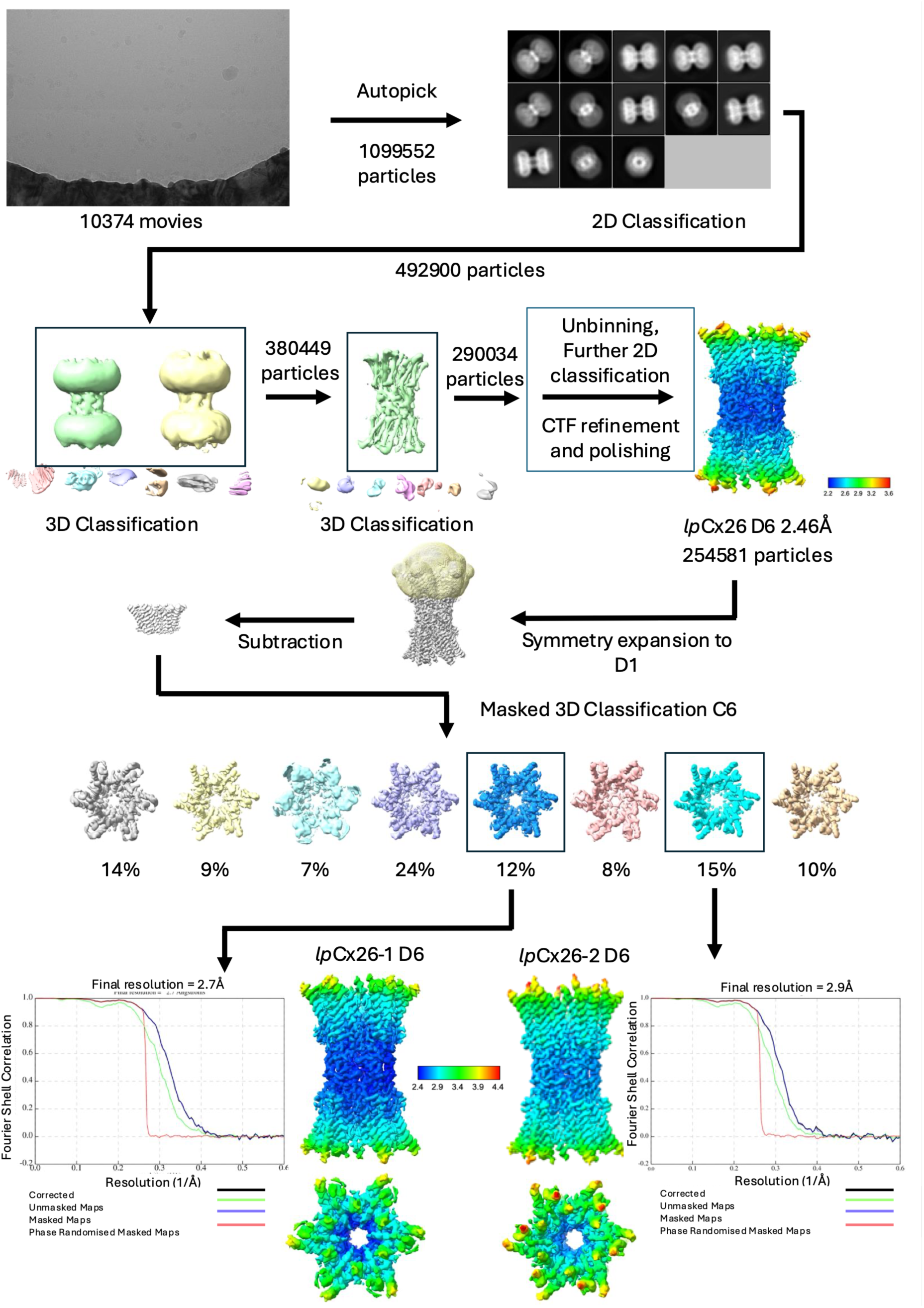
Workflow for processing of cryo-EM data for *lp*Cx26 purified in GDN. From the maps derived from the focussed classification *lp*Cx26-GDN-1 and *lp*Cx26-GDN-2 were chosen based on the clarity of the N-terminus and TM3 respectively.

**Supplementary Figure 5:**
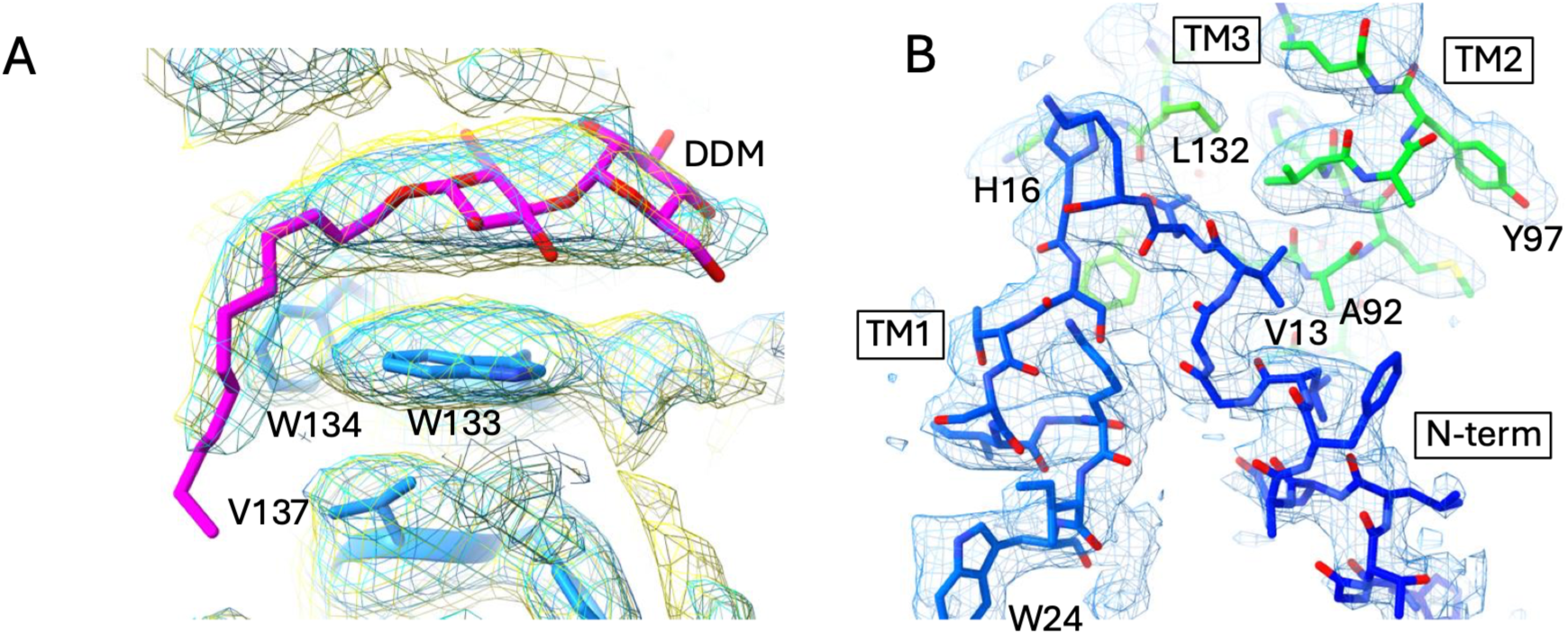
Density associated with *lp*Cx26 exchanged into GDN A) Comparison of density around the putative DDM for *lp*Cx26-GDN-1 (blue), *lp*Cx26-GDN-1 (cyan) and *lp*Cx26-DDM (yellow). B) Density associated with *lp*Cx26-GDN-1 around the N-terminus/TM1 loop.

**Supplementary Figure 6:**
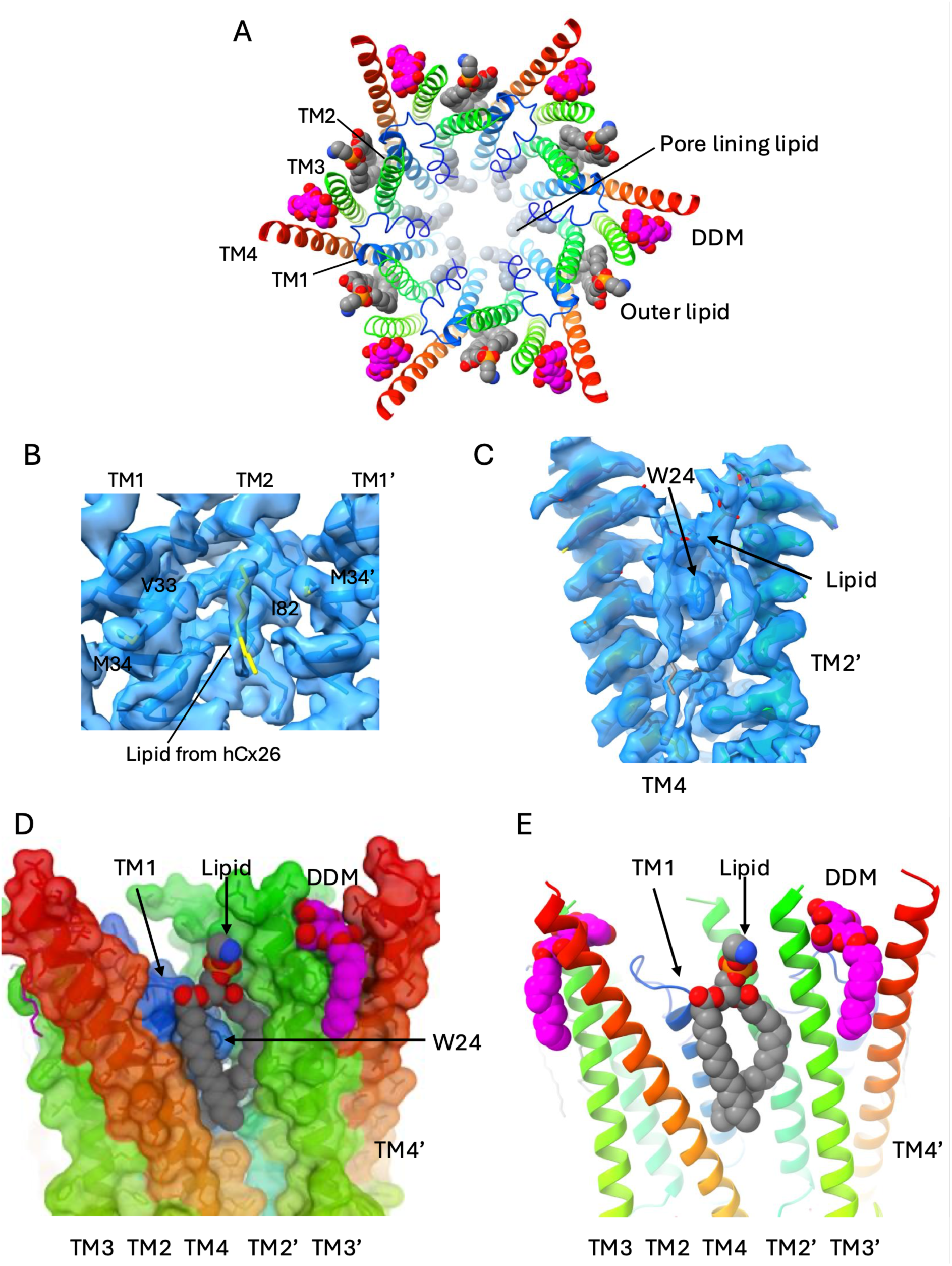
Lipid binding to *lp*Cx26. A) View looking into the pore from the cytoplasmic side. The protein has been coloured from blue to red as shown for Supplementary Figure 1. The lipids and DDM are depicted as spheres with grey carbon atoms for the lipid and magenta carbon atoms for the DDM. B) Density (*lp*Cx26-GDN-1) associated with the lipid located in a crevice between TM2, the N-terminus and TM1 from two neighbouring subunits in the pore of the protein. Residues of *lp*Cx26 are shown with blue carbon atoms. The lipid molecule in yellow is from hCx26 (PDB 8QA0). C) Density (*lp*Cx26-GDN-1) associated with the lipid molecule on the outer face of the pore as seen in A. D) View of the lipid as shown in C. The lipid is sandwiched between TM1 and TM4 of one molecule and TM2 and TM3 of the next with DDM on the other side of TM3. The surface representation of the protein has been coloured as in A. E) as D but with the surface removed.

**Supplementary Figure 7:**
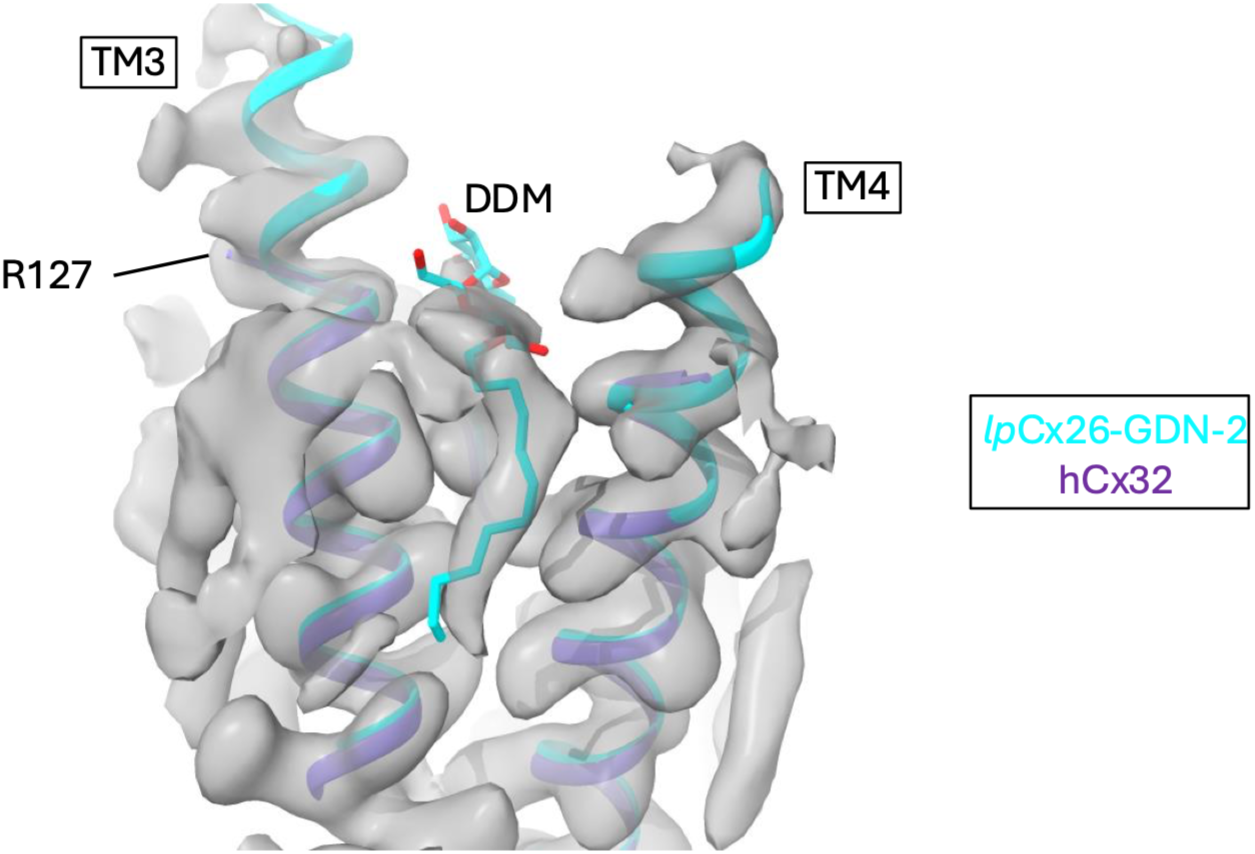
Comparison to Cx32 GJCs. Comparison of lpCx26 (cyan) with the structure of hCx32 (purple, PDB 9QNF) and associated map (EMD-53245). While TM3 of the deposited structure is only built from Arg127, the helical density extends further as seen for *lp*Cx26. Density is also apparent in the map between TM3 and TM4 in the same position as DDM from *lp*Cx26.

**Supplementary Figure 8:**
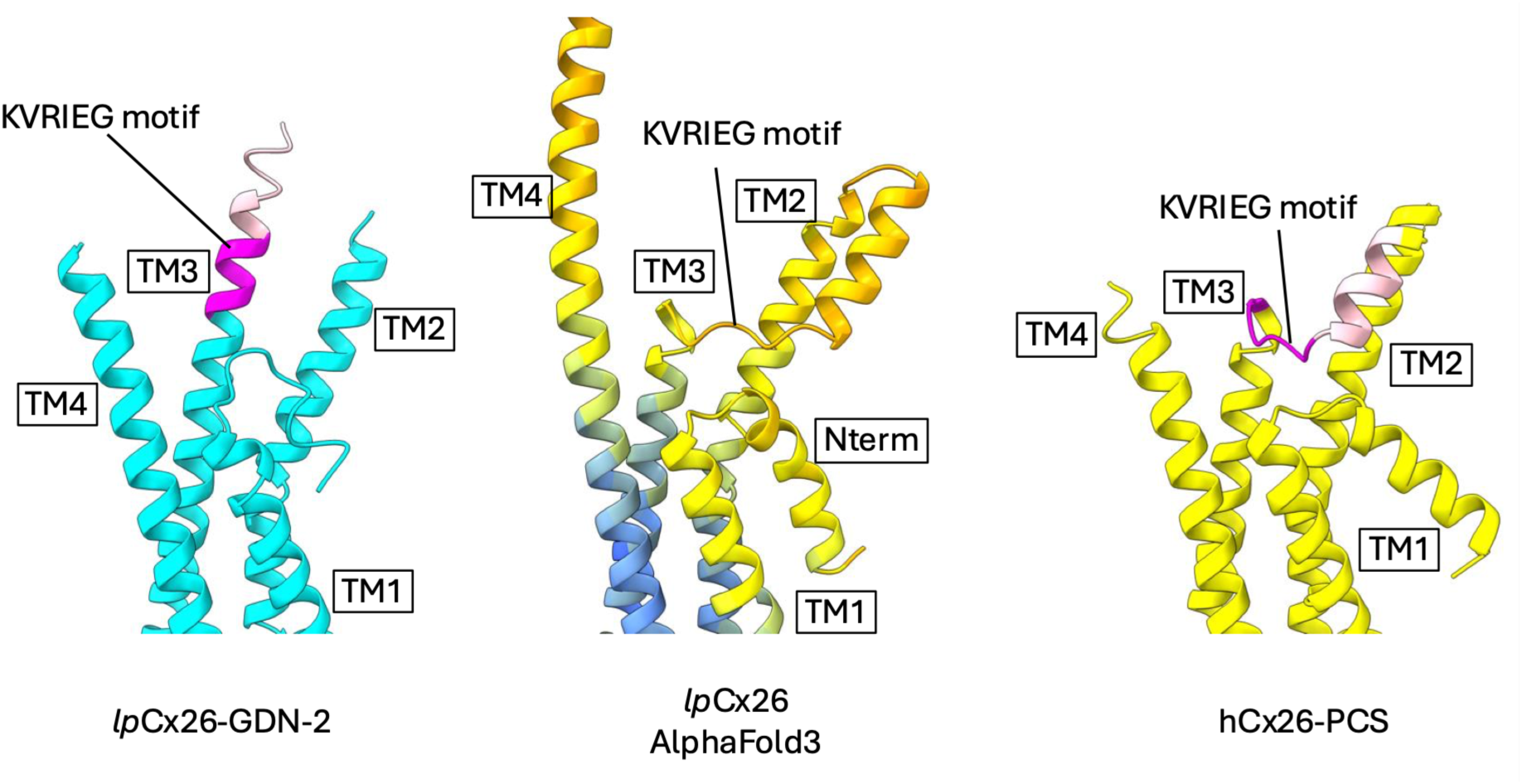
Comparison of *lp*Cx26 to AlphaFold3 model. Side by side views of the superposed structures of the A subunit of lpCx26-GDN-2 (left), an AlphaFold3 prediction of *lp*Cx26 (middle) and the structure of hCx26 pore constricted state elucidated through classification focussed on single subunits (right; PDB 8QA2). The lpCx26-GDN-2 and hCx26-PCS structures have been coloured cyan and yellow respectively, with the residues of the KVRIEG motif coloured magenta and residues 117-124 hot pink. The AlphaFold3 prediction of lpCx26 has been coloured according to the AlphaFold colouring palette in ChimeraX where the warmer colours denote less confidence. While the residues around the KVRIEG motif are not predicted with high confidence, the predominant predicted conformation for Cx26 positions the KVRIEG motif in a similar conformation to that seen in the hCx26 pore constricted state.

